# SIRT5-mediated GLS and GDH desuccinylation attenuates autophagy in MAC-T cells induced by ammonia

**DOI:** 10.1101/2024.08.26.609685

**Authors:** Hanlin Yang, Shikai Gao, Guangyang Lu, Junhui He, Jinru Dong, Xinyi Zhang, Luya Liu, Kai Zhong, Guangming Zha, Liqiang Han, Shuang Guo, Heping Li, Yueying Wang

## Abstract

Our previous research revealed that NH_3_ regulated autophagy dependent on SIRT5 in MAC-T cells. Interestingly, SIRT5 reduced the content of NH_3_ and glutamate by inhibiting GLS activity, ADP/ATP value also declined. In this study, SIRT5 interacted with endogenous GLS and GDH, and had no effect on endogenous GLS and GDH expression. SIRT5 declined significantly the succinylation levels of GLS and GDH, and further reduced the enzymatic activity of GLS and GDH. SIRT5 declined the glutamine metabolism, which attenuated ammonia release in MAC-T cells, accompanying with cellular autophagy decline, reducing the formation of autophagosome. Deletion of SIRT5 increased the content of NH_3_ and glutamate, as well as promotes autophagy, which could be alleviated by SIRT5 overexpression. SIRT5 KO was associated with increased succinylation and activity of GLS and GDH, as well as autophagy response in MAC-T cells. Furthermore, SIRT5 promoted the maintenance of mitochondria homeostasis. Mechanistically, SIRT5 modulated the succinylation levels and enzymatic activities of GLS and GDH in mitochondria and promoted the maintenance of mitochondria homeostasis, further attenuating ammonia-stimulated autophagy in MAC-T cells.

**Highlights:**

- SIRT5 catalyzed lysine desuccinylation of GLS and GDH.
- GLS and GDH enzymatic activity were enhanced by lysine succinylation.
- GLS and GDH were required for SIRT5 to regulate ammonia-induced cellular autophagy.
- SIRT5 promoted the maintenance of mitochondrial homeostasis

## Introduction

Ammonia (NH_3_), as one of the harmful gases in livestock houses, affected the health and growth of livestock, and reduced their production performance [1]. In addition, NH_3_ could change pH, electrolyte balance and metabolic changes, inducing various transform in many cell types [2]. In vitro, ammonia had a dual role in autophagy, it played an inducer at lower concentrations and an inhibitor at high concentrations [3]. Ammonia at high concentrations (>20 mM) enhanced pH in lysosome and increased water influx, resulting in inhibition of substrate degradation and lysosome swelling, and further destroying lysosomal function [4, 5]. In contrast, ammonia strongly promoted autophagy at lower concentrations (0.2∼10 mM) [6–12]. Ammonia derived from glutamine catabolism also strongly induced autophagy in cancer cells [13]. Apoptosis and autophagy exert key role in maintaining the quantity and ability of mammary epithelial cells, and are key factors affecting the lactation performance of dairy cows. Our previous research found that NH_3_ regulated the autophagy in cow mammary epithelial cells through PI3K/Akt/mTOR signaling pathway [14].

Sirtuin 5 (SIRT5) was an important regulatory factor in maintaining cellular homeostasis, which emerged to be an important desuccinylase enzyme, desuccinylating more than half of proteins in mitochondria and controlling ammonia-induced autophagy in tumor cells [15–23]. In addition, we found that SIRT5 inhibited the autophagy in bovine mammary epithelial cells through PI3K/Akt/mTOR signal, involving in NH_3_-induced autophagy dependent on SIRT5 [24]. Moreover, there were many studies explaining the dual role of SIRT5 for autophagy. SIRT5 promoted autophagy in colorectal cancer, where its overexpression was associated with low survival [20]. SIRT5 promoted autophagy in gastric cancer cells through the AMPK/mTOR pathway, while SIRT5 expression was suppressed in gastric cancer tissues [25]. In breast cancer and mouse myogenic cells, silencing SIRT5 increased ammonia-induced autophagy by controlling glutamine metabolism and mitophagy [12]. The glutaminolysis in mitochondria was catalyzed by glutaminase (GLS) with deamination of glutamine, by which produced ammonia and glutamate. Glutamate was converted to α-ketoglutarate through oxidative deamination by glutamate dehydrogenase (GDH) in mitochondria, releasing NH_4_^+^ [26, 27]. SIRT5 had been shown to exhibit weaker deacetylase activity, more commonly demalonylase, desuccinylase, and deglutarylase activities, participating in several metabolic processes of mitochondria [18, 28–31]. SIRT5 downregulated the succinylation level of GLS and inhibited ammonia-induced autophagy in tumor cells [12]. Our previous studies also found that SIRT5 reduced NH_3_-induced autophagy in bovine mammary epithelial cells, but little is known about the specific molecular mechanism.

Interestingly, our recent study found that SIRT5 reduced the content of NH_3_ and glutamate in bovine mammary epithelial cells by inhibiting GLS activity, and declined the ratio of ADP/ATP [24]. Therefore, we hypothesized that SIRT5-mediated GLS and GDH desuccinylation controlled ammonia production, further regulated the autophagy of mammary epithelial cells. To verify this point, we constructed SIRT5 overexpression or knockout cell lines. Additionally, ammonium chloride, and some inhibitors for SIRT5, GLS, and autophagy were also used to treat cells. The interaction of SIRT5 with GLS and GDH were identified using immunoprecipitation techniques, and the succinylation level and enzyme activity of GLS and GDH were also determined. The results verified that SIRT5 interacted with endogenous GLS and GDH, and had no effect on the expression of endogenous GLS and GDH. Expectedly, we discovered that SIRT5 obviously decreased the succinylation levels of GLS and GDH, and further reduced the enzymatic activity of GLS and GDH. Next, the content of ammonia and glutamate, as well as the related autophagy markers were measured, the results demonstrated that SIRT5 declined the glutamine metabolism and ammonia release in MAC-T cells, accompanying with cellular autophagy decline and less autophagosome formation. Interestingly, SIRT5 enhanced the content of ATP and promoted the maintenance of mitochondria homeostasis. Altogether, SIRT5 reduced ammonia release by modulating the succinylation levels and enzymatic activities of GLS and GDH in mitochondria, and further promoted the maintenance of mitochondria homeostasis, accompanying with attenuating ammonia-stimulated autophagy in cow mammary epithelial cells. In this research, we revealed that GLS and GDH were two physiological substrates of SIRT5, which relied on desuccinylation to reduce ammonia production by inhibiting the enzymatic activity of GLS and GDH.

## Materials and methods

### Generation of SIRT5 OE and KO bovine mammary epithelial cell lines

Bovine mammary epithelial cells (MAC-T cells, preserved in our laboratory) were cultured according to our previous method [14, 24]. To generate SIRT5 overexpression cells (SIRT5 OE), bovine SIRT5 gene (NM_001034295) was subcloned into a pPB-EF1α eukaryotic expression vector (Haixing Biotechnology Co., Ltd). The vectors with SIRT5 (pPB-EF1α-SIRT5 vector) or without SIRT5 (pPB-EF1α vector), which contained non-fused EGFP fluorescent tags and resisted to puromycin, were electrotransfected into MAC-T cells using the Neon transfection system (Neon™ transfector, Cat# MPK5000) according to the product manual (Thermo Fisher Scientific Inc.). The empty vector pPB-EF1α without the insert (SIRT5 cDNA) was used as the control. At 48 h post-transfection, the generation of SIRT5 OE cell line was achieved by selecting with 5.0 μg/mL puromycin.

CRISPR-Cas9 mediated the excision of SIRT5 gene was obtained with CRISPR-Cas9 RNP (supplied with Haixing Biotechnology Co., Ltd) including expression element for hSpCas9 and chimeric lead RNA. To guide exon 4∼exon 9 of SIRT5 gene, two target RNA sequence of AGCGTGCTTTCCCGAGACAGCGG and GCGGGTGACGGAGTTGTGTGTGG were selected on the http://crispr.mit.edu website. Vector including the target RNA sequence was electrotransfected into cells using the Neon transfection system according to the product manual (Thermo Fisher Scientific Inc.). After 48 h, single colonies were moved to 96-well culture plates. To measure the exist of indels in SIRT5 guided colony, a Quick-DNA Miniprep kit (Zymo Research, USA, CA) was used to segregate genomic DNA, and 2×Taq Master Mix (Dye Plus, Vazyme, P112) with exon flanking primers was obtained for PCR amplification. Forward: 5’-AGTGGGACGGAG CATTTGTT-3’; Reverse: 5’-CTGACTTAGGTAATGACAAGATGCT-3’. Vectors were collected from 8-10 single clones to be sequenced by Sanger sequencing (GENEWIZ, China). Colonies with mutations in both alleles were chosen for subsequent tests. SIRT5 knockout (KO) cell line was obtained according to the above steps, and cultivated under the same conditions as the parental cells. Then, we used PCR and western blotting method to verify SIRT5 expression in SIRT5 OE and KO MAC-T cell lines. RT-PCR method was conducted according to our previous literatures [14, 24]. Glyceraldehyde-3-phosphate dehydrogenase (GAPDH) was chosen as internal reference genes. The primers of SIRT5 and GAPDH as following. GAPDH (NM_001034034.2): (upstream, 5’-3’) CCATGTTTGTGATGGGCGTG, (downstream, 5’-3’) GCAGGGATGATATTCTGGGCA; SIRT5 (NM_001034295.2): (upstream, 5’-3’) TTGTGGAGTTGTGGCTGAGA, (downstream, 5’-3’) GTCCCCACCACTAGACACAG. The details of western blotting method were described later.

### Cell treatment

Ammonium chloride (NH_4_Cl, Sigma-Aldrich, Shanghai, A43A4899) as a donor of NH_3_ was diluted in deionized water and adjusted to an ultimate concentration of 10 mM. MC3482 (SIRT5 inhibitor) from MedChemExpress (Monmouth Junction, NJ, USA) was diluted in deionized DMSO and adjusted to an ultimate concentration of 20 μM. BPTES (GLS inhibitor) and BafA1 (bafilomycin A1, autophagy inhibitor) were purchased from MedChemExpress LLC (Shanghai). The MAC-T cell and SIRT5 KO, SIRT5 OE cell lines were cleaned twice using phosphate-buffered saline and then cultivated in medium containing the indicated agents. The MAC-T cells were randomized into the seven experimental groups: (1) Control (CT), (2) NH_4_Cl (NC), (3) MC3482 (MC), (4) MC3482+NH_4_Cl (MCN), (5) BPTES (BP), (6) MC3482+BPTES (MCBP), (7) BafA1 (BA). The SIRT5 OE cell lines were randomized into the three experimental groups: (1) SIRT5 OE (SO), (2) SIRT5 OE+NH_4_Cl (SON), and (3) SIRT5 OE+BPTES (SOBP). The SIRT5 KO cell lines were randomized into the four experimental groups: (1) SIRT5 KO (SK), (2) SIRT5 KO+NH_4_Cl (SKN), (3) SIRT5 KO+BPTES (SKBP), and (4) SIRT5 KO+NH_4_Cl+BafA1 (SKNBA). Each of the above groups contained three independent repeats. In CT, SO, and SK groups, the MAC-T, SIRT5 OE, and SIRT5 KO cells were cultivated for 12 h in basal medium, respectively. In the NC, SON, SKN groups, the MAC-T, SIRT5 OE, and SIRT5 KO cells were treated with 10 mM NH_4_Cl for 12 h, respectively. In the MC, BP, and BA group, MAC-T cells were incubated with 20 μM MC3482, or BPTES (0.12 μM), or BafA1 (50 nM) for 12 h, respectively. In the MCN and MCBP group, MAC-T cells were incubated with MC3482 (20 μM) for 1 h, and then the cells were exposed to 10 mM NH_4_Cl or BPTES (0.12 μM) for 12 h. In SOBP, SKBP groups, SIRT5 OE cells and SIRT5 KO cells were cultivated in medium containing BPTES (0.12 μM) for 1 h, and then the cells were exposed to 10 mM NH_4_Cl for 12 h. In SKNBA group, SIRT5 KO cells were treated with BafA1 (50 nM) for 1 h, and then the cells were exposed to 10 mM NH_4_Cl for 12 h.

### Immunoprecipitation assay

For detection of protein-protein interactions, cells were lysed with 500 μL of specific immunoprecipitation lysis solution for 30 min with orbital vibrating on ice. Then, lysates were selected and moved into cold microfuge tubes, at the centrifugal force of 12, 000 g for 10 min at 4℃. For co-immunoprecipitation (Co-IP) of immune complexes, cell lysate was first pre-cleared by inoculation with 50 μL of protein A/G magnetic beads (Thermo Fisher Scientific Inc.) and rabbit anti-IgG (to decline most of unspecific protein interactions) for 30 min at 4°C with lightly vertical mixing (10 RPM). The beads were gathered with a magnet, and the lysates were moved into a new, cold microfuge tube.

Next, 5 μL GLS, SIRT5 or GDH antibodies were replenished into the tube, respectively. The whole volume was increased to 500 μL. The samples were inoculated at 4°C for overnight with lightly vertical mixing (10 RPM) to form immune complex. The magnetic beads of protein A/G were wetted and washed triple using PBST, and then the incubated lysates were bound to the magnetic beads and held in a rotator at 25°C for 6 h. After binding, the lysates were wetted and washed three times at 99°C for 10 min with PBST and 70 μL 1×SDS loading buffer. The beads were collected, and the lysates were moved into a new, cold microfuge tube. Samples were then electrophoresed on a 10 % SDS polyacrylamide gel and further immunoblotted with designated antibodies. Here, IgG was selected as a negative control. ImageJ software was adopted to analyze grayscale values, calculate relative protein expression levels, and perform statistical analysis.

Mouse anti-SIRT5 monoclonal antibody (Cat No:67257-1-Ig; 1:1000), rabbit anti-GLS polyclonal antibody (Cat No:12855-1-AP; 1:2000), rabbit anti-β-actin recombinant antibody (Cat No:81115-1-RR; 1:5000), rabbit IgG control polyclonal antibody (Cat No:30000-0-AP; 1:2000) and goat anti-rabbit IgG (H+L) (Cat No: SA00001-2; 1:5,000) were purchased from Proteintech Group, Inc. Rabbit anti-SIRT5 monoclonal antibody (Cat No: ab259967; 1:1000) and rabbit monoclonal anti-GDH antibody (Cat No: ab166618; 1:2000) were purchased from Abcam (Shanghai) trading Co., Ltd. Rabbit anti-succinylated lysine polyclonal antibody (Cat No: 3089; 1:2000) was brought from Wuhan DIA AN biotechnology Co., Ltd.

### Biochemical assays

The cells were lysed, and centrifuged. Then, the supernatant was collected. The BCA protein assay kit (Abcam, Shanghai, ab102536), ammonia assay kit (Sigma-Aldrich, Shanghai, AA0100), glutamate measurement kit (Nanjing Jiancheng Bioengineering Institute, China, A074-1-1), ATP assay kit (Beyotime Biotechnology, China, S0026), glutaminase test kit (Nanjing Jiancheng Bioengineering Institute, China, A124-1-1), and glutamate dehydrogenase test kit (Nanjing Jiancheng Bioengineering Institute, China, A125-1-1) were used to determine the protein concentration, the content of ammonia, glutamate and ATP, as well as GLS and GDH enzymatic activity. The above kits were operated according to the manufacturer’s specifications. All assays were executed in triplicates.

### Western blotting

Western blotting was used to analyze the expression of SIRT5 and designated autophagy proteins. The above treated cells were cleared twice with cold PBS, the BCA method was applied to measure the concentration of protein. Next, 5×SDS loading buffer was increased to the extracted proteins, which were then denatured at 99°C for 10 min. The proteins (24 μg/lane) were separated with 10 % SDS-PAGE gels, and then moved onto polyvinylidene fluoride (PVDF) membranes (EMD Millipore) that were sealed with 5 % skim milk for 1 h. The membranes were sequentially incubated with the specific first antibodies at 4°C for overnight. The membranes were cleared with TBST for 30 min and inoculated with goat anti-rabbit IgG (H+L) antibody (Cat No: SA00001-2; 1:5,000; purchased from Proteintech Group, Inc.) at room temperature for 2 h. Then, the membranes were washed with TBST for 30 min again. Finally, the visualized protein bands were detected under a developer using enhanced chemiluminescence reagent (EMD Millipore), and ImageJ software 1.4.3.67 (National Institutes of Health) was adopted to evaluate the grayscale values of the protein bands, and calculate the relative expression of proteins in each group.

The corresponding primary antibodies used as following: rabbit anti-SIRT5 monoclonal antibody (Cat No: ab259967; 1:1000) was purchased from Abcam (Shanghai) trading Co., Ltd. Rabbit anti-Beclin1 polyclonal antibody (Cat No: 11306-1-AP; 1:2000), rabbit anti-LC3B polyclonal antibody (Cat No: 18725-1-AP; 1:1000), and rabbit anti-p62/SQSTM1 polyclonal antibody (Cat No: 18420-1-AP; 1:5000) were bought from Proteintech Group, Inc.

### Immunofluorescence staining

To identify the formation of autophagosomes through LC3B puncta, immunofluorescence staining was performed on the cells. Briefly, MAC-T cells were dealt with or without NH_4_Cl, and SIRT5 KO cells were treated with NH_4_Cl or NH_4_Cl and BafA1 together. Cells achieving a cell density of 50 % were cultured in a 24-well plate. 4 % paraformaldehyde was used to fix the cells, at room temperature for 30 min and rinsed with PBS. Then, 0.2 % Triton X-100 was added for 10 min at room temperature, and the cells were rinsed with PBS for 30 min. Next, 10 % fetal bovine serum (FBS) was added to block for 1 h at room temperature, after which the anti-LC3B antibody was added to inoculation at 4°C for overnight. The secondary fluorescent antibody was inoculated at room temperature with protection from light for 1 h, and the cells were cleared in PBS. 4, 6-diamidino-2-phenylindole (DAPI) was used to performe nuclear staining. Finally, anti-fluorescence quenching sealer was added and the film was sealed and stored, and the stained LC3B positive cells was observed with a laser confocal fluorescence microscope after overnight stabilization. The effect of SIRT5 on the formation of autophagosomes was evaluated with the number of autophagosomes, which was calculated based on the number of green fluorescent aggregated spots from three views in each sample. Rabbit polyclonal to LC3B-autophagosome marker (Cat No: ab48394; 1:500; Abcam (Shanghai) trading Co., Ltd) goat anti-rabbit IgG (H+L) cross-adsorbed secondary antibody, Alexa Fluor™ 488 (Cat No: A-11008; 1:500; Cell signaling technology, Inc.) were used.

### Mitochondrial staining

Mito-Tracker Red staining is a method for labeling mitochondria, which binds to deoxyribonucleotides inside the mitochondria to stain the mitochondria of living cells. Mito-Tracker Red CMXRos (Cat No: C1049B) was used in this study, and bought from Shanghai Beyotime Biotechnology, Inc. Mito Tracker Red CMXRos is a kind of cell permeable agent derivating from X-rosamine (Chloroethyl-X-rosamine, abbreviated as CMXRos) that can specifically label biologically active mitochondria in cells. As the aggregation of Mito Tracker Red CMXRos in mitochondria depends on the membrane potential of mitochondria, it can only stain live cells and detect mitochondrial membrane potential. Using Mito-Tracker Red-stained MAC-T cells (CT), SIRT5 OE, and SIRT5 KO cells, mitochondria was observed under confocal laser microscopy, and further analyzed the effects of SIRT5 on the morphology and quantity of mitochondria, and mitochondrial activity.

### Statistical analysis

The quantitative data were displayed as the mean±standard deviation. GraphPad Prism software (version 6.01, GraphPad, Dotmatics) was selected for statistical analysis. Differences among groups were calculated using one-way ANOVA followed by post-hoc sidak’s correction. P<0.05 was indicated as a statistically significant difference.

## Results

### Generation of SIRT5 OE and KO MAC-T cell lines

The fluorescence intensity in cells was evaluated under a fluorescence microscope to evaluate the successful transfection. As seen in Figure 1A, no fluorescence existed in non-transfected MAC-T cells. A large amount of green fluorescence was observed in cells transfected with the pPB-EF1α vector and the pPB-EF1α-SIRT5 vector (Figure 1B, C). Meanwhile, the PCR and western blotting results verified that SIRT5 expression at the mRNA and protein level in SIRT5 KO cells was almost undetectable (Fig. 2A, B), while SIRT5 expression was promoted at the mRNA and protein level in SIRT5 OE cells compared to MAC-T cells, and the difference significant was obvious (Fig. 2C, D). Combined with the observation of fluorescence microscopy, the results showed that the SIRT5 OE and SIRT5 KO cell lines were successfully generated.

**Figure 1.**
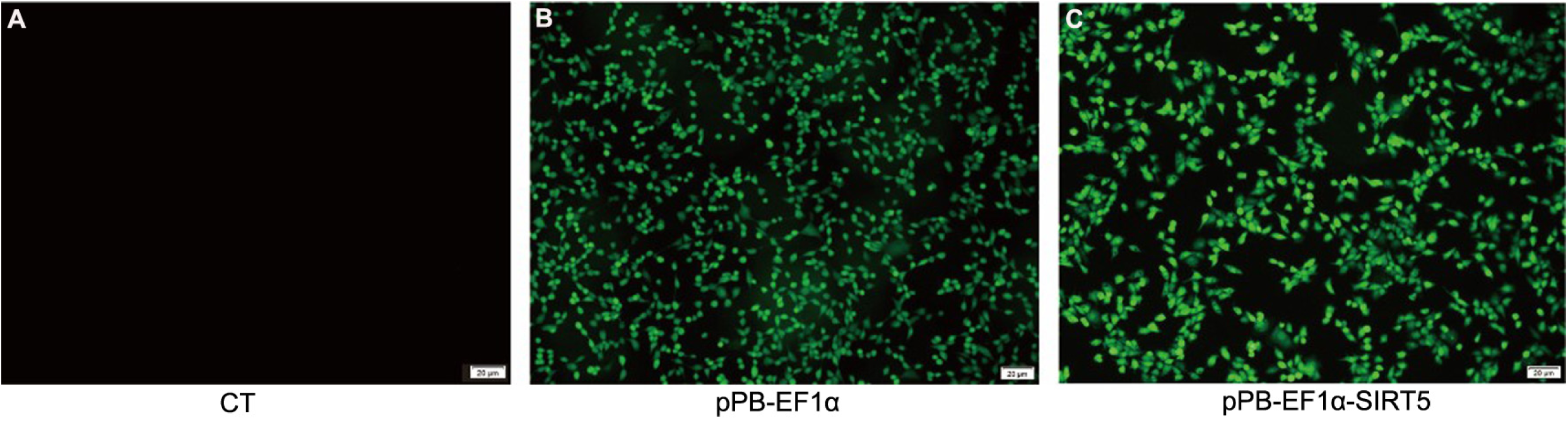
Observation of green fluorescence in MAC-T cells transfected with pPB-EF1α or pPB-EF1α-SIRT5 vectors. Green fluorescence was observed clearly under the fluorescence microscopy in the transfected MAC-T cells (B, pPB-EF1α; C, pPB-EF1α-SIRT5), but no green fluorescence in non-transfected MAC-T cells (A, CT).

**Figure 2.**
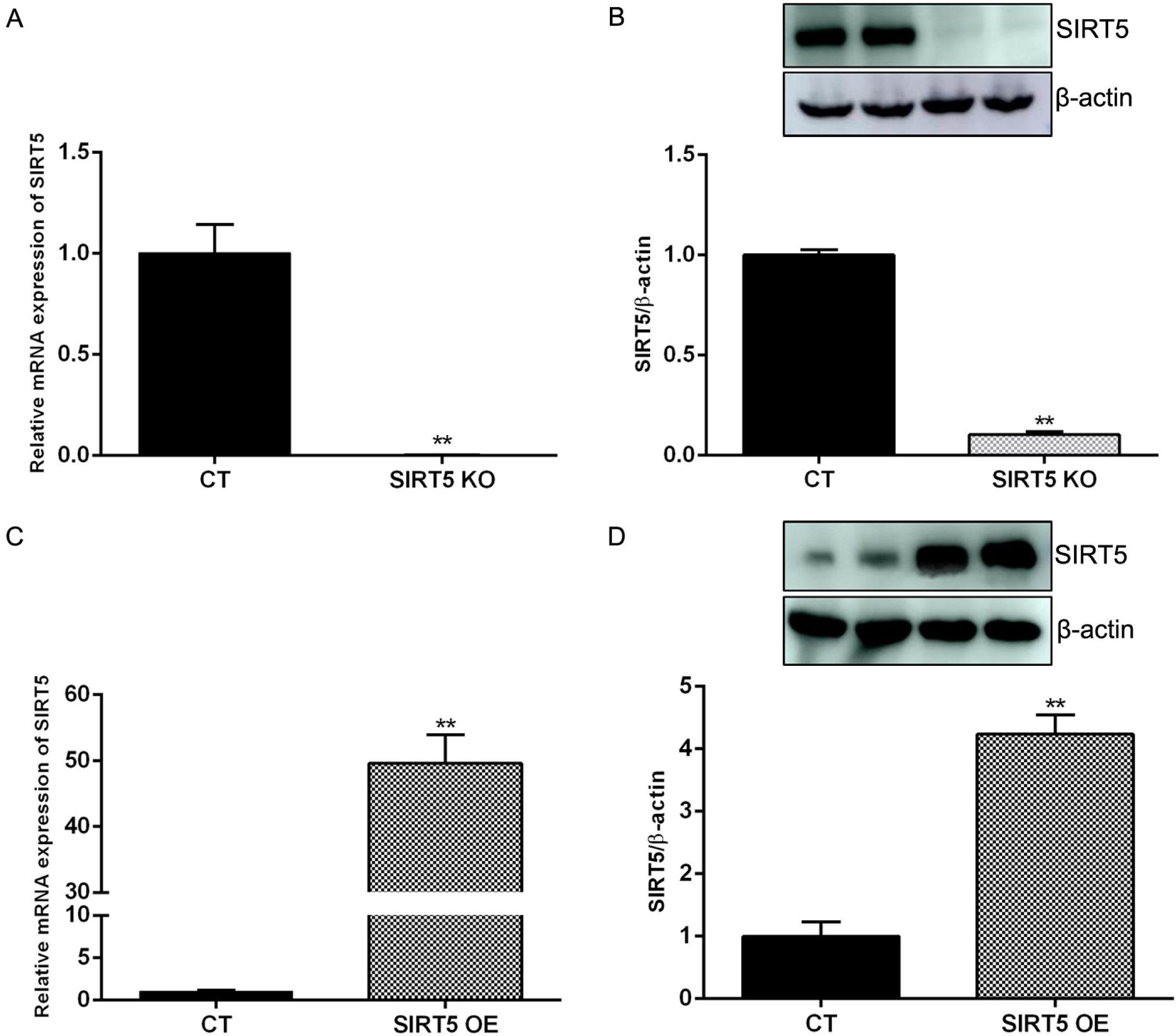
SIRT5 expression in SIRT5 KO and OE cell lines. (A, B) The mRNA and protein expression of SIRT5 in SIRT5 KO cells. (C, D) The mRNA and protein expression of SIRT5 in SIRT5 OE cells. Note: ** P < 0.01 vs. MAC-T cells (CT group).

### SIRT5 catalyzed lysine desuccinylation of GLS and GDH

SIRT5 in mitochondria had a strong lysine desuccinylation capacity, involving in several processes of mitochondrial metabolism [18, 28–31]. Interestingly, our previous study found that SIRT5 reduced the content of NH_3_ and glutamate in bovine mammary epithelial cells by inhibiting GLS activity [24]. As list in Figure 3, SIRT5 interacted with endogenous GLS or GDH. Strangely, SIRT5 had no effect on the expression of endogenous GLS or GDH (Figure 4). Compared with MAC-T cells, the protein succinylation levels were down-regulated with statistical difference in SIRT5 OE cells and significant up-regulated in SIRT5 KO cells (Figure 5). These demonstrated that SIRT5 declined the protein succinylation level of cells. Interestingly, we also found that the GLS or GDH succinylation level was significantly down-regulated in SIRT5 OE cells, whereas the GLS or GDH succinylation level was obviously up-regulated in SIRT5 KO cells (Figure 6). Taken together, SIRT5 interacted with endogenous GLS or GDH, and further catalyzed lysine desuccinylation of GLS and GDH.

**Figure 3.**
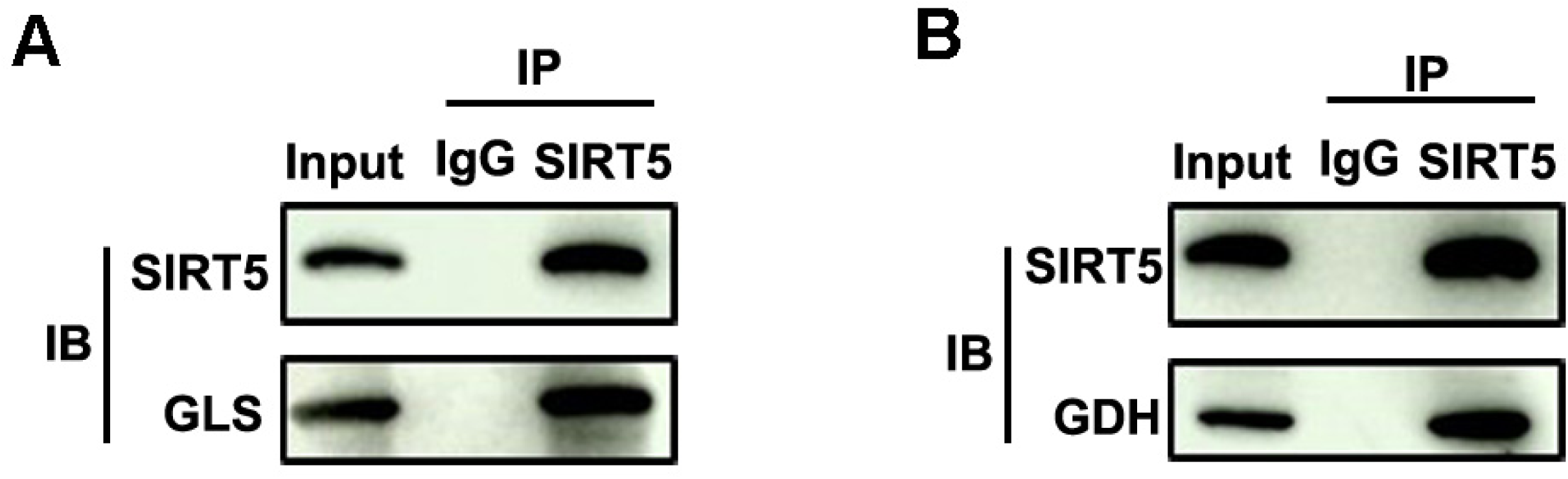
SIRT5 interacted with endogenous GLS or GDH. A. Endogenous GLS interacts with SIRT5. SIRT5 protein in MAC-T cells was purified by IP with an anti-GLS antibody, followed by western blotting to detect SIRT5 with an anti-SIRT5 antibody. B. Endogenous GDH interacts with SIRT5. SIRT5 protein in MAC-T cells was purified by IP with an anti-GDH antibody, followed by western blotting to detect SIRT5 with an anti-SIRT5 antibody.

**Figure 4.**
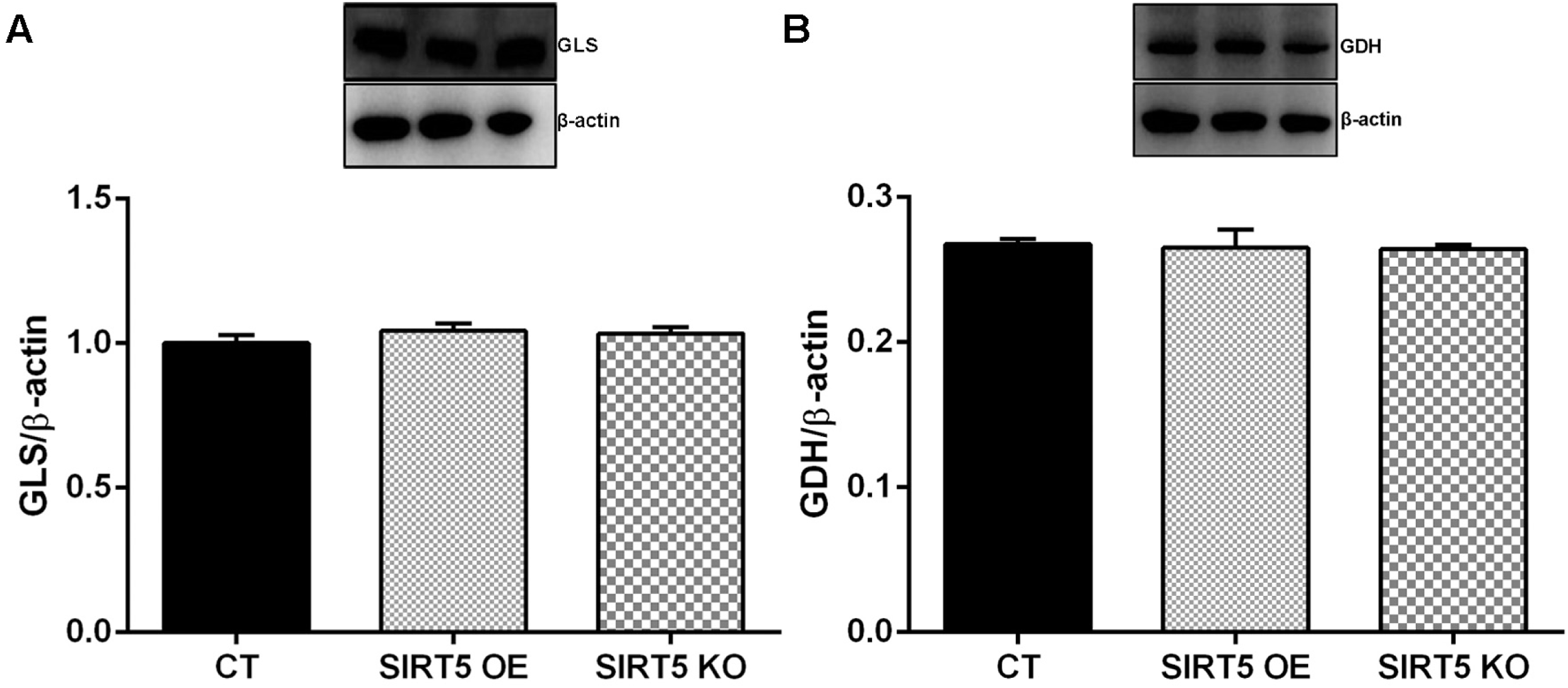
SIRT5 did not change the expression of GLS and GDH. A. GLS expression detected by western blotting. B. GDH expression detected by western blotting. β-actin is used as loading control. MAC-T cells (CT), SIRT5 OE and SIRT5 KO cells were cultured in base medium, respectively.

**Figure 5.**
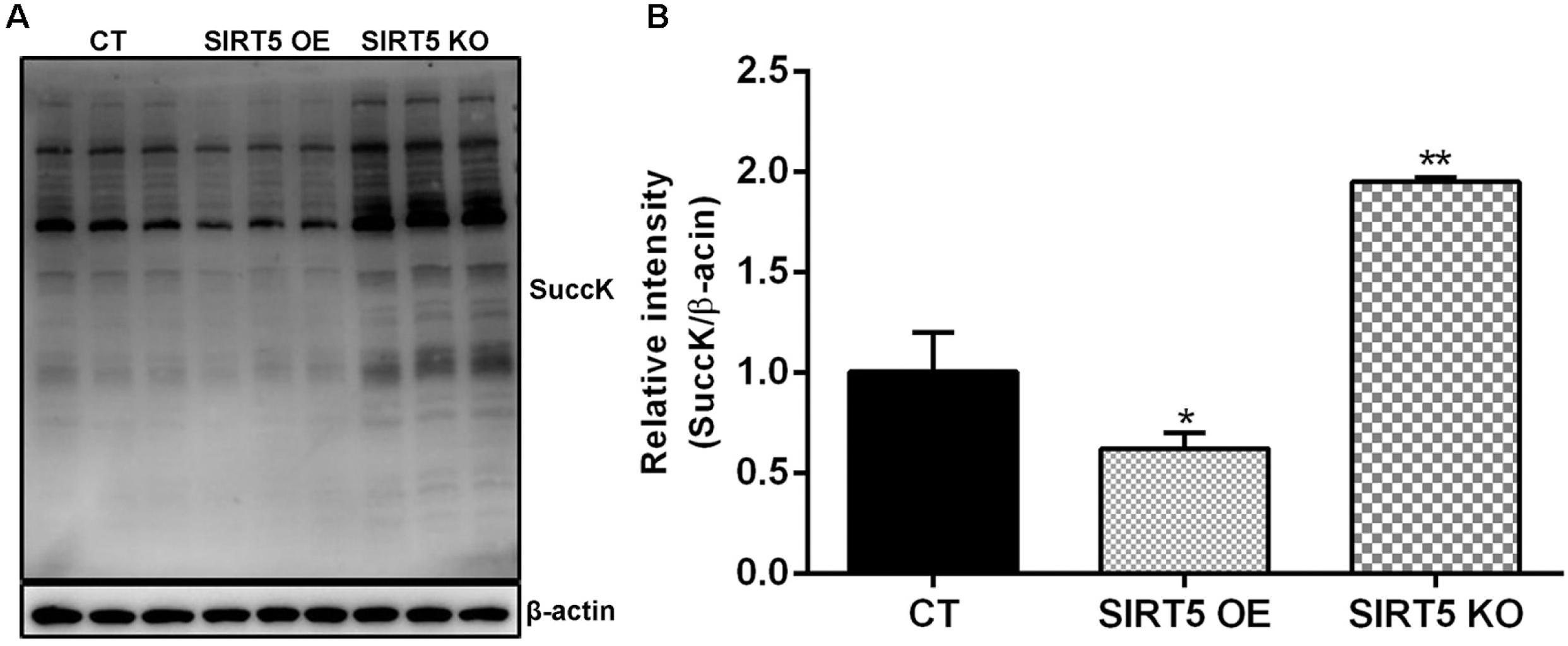
SIRT5 decreased protein succinylation in whole cell lysates. A. Immunoprecipitation and western blotting were performed to assess succinylation levels using succinyl-lysine-specific antibodies in whole cell lysates. B. Relative intensity of SuccK/β-actin. In stable MAC-T, SIRT5 OE, and SIRT5 KO cells, the total proteins were purified by IP with beads and western blotting to detect their lysine succinylation level. Note: * P < 0.05, ** P < 0.01 vs. MAC-T cells (CT group).

**Figure 6.**
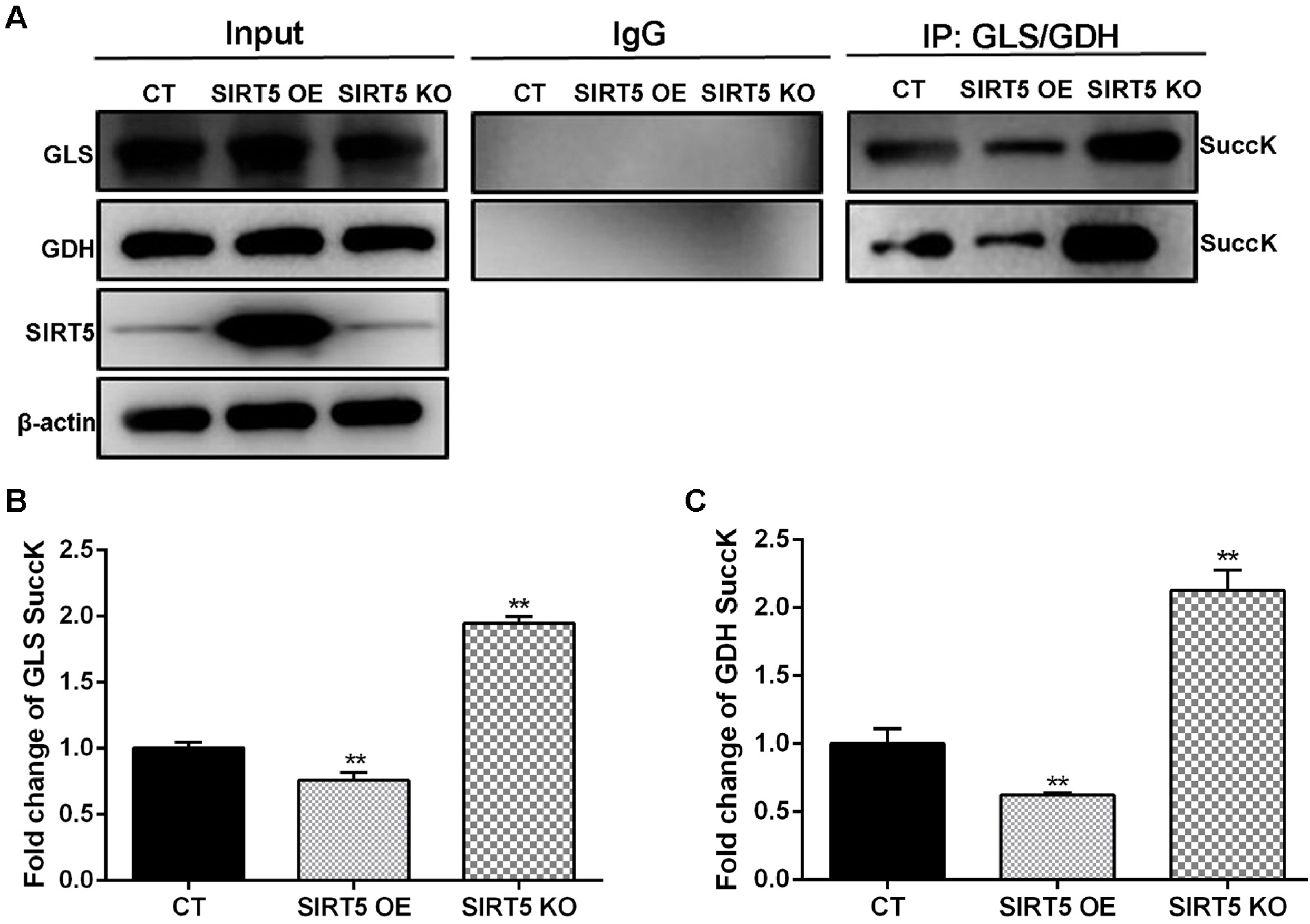
SIRT5 decreased the GLS and GDH succinylation. A. Immunoprecipitation and western blotting were performed to assess succinylation levels of GLS and GDH using succinyl-lysine-specific antibodies. B. Fold change of GLS SuccK. C. Fold change of GDH SuccK. In stable MAC-T, SIRT5 OE, and SIRT5 KO cells, the GLS and GDH protein were purified by IP with beads and western blotting to detect their succinylation level. Note: ** P < 0.01 vs. MAC-T cells (CT group).

### GLS and GDH enzymatic activity were enhanced by lysine succinylation

To determine the functional consequence of lysine desuccinylation of GLS and GDH catalyzed by SIRT5, we detected the GLS and GDH enzymatic activity. In SIRT5 OE cells, GLS and GDH enzymatic activity were lower than those in MAC-T cells (Figure 7A, 7B). Similar changes existed in the content of NH_3_ and glutamate (Figure 7C, 7D). Correspondingly, with NH_4_Cl treatment, SIRT5 also reduced GLS and GDH enzymatic activity, resulting in the induction of NH_3_ and glutamate content, alleviating glutamine metabolism (Figure 8). In contrast, SIRT5 KO cells led to a profound increase in GLS and GDH enzymatic activity (Figure 7), which were associated with increased succinylation level of GLS and GDH (Figure 6). Meanwhile, compared with the CT group, inhibiting SIRT5 (MC3482, a specific SIRT5 inhibitor) improved obviously GLS and GDH enzymatic activity (Figure 7A, 7B), accompanying with obvious reduction in the content NH_3_ and glutamate (Figure 7C, 7D). In addition, compared to SIRT5 OE or KO cells, we observed that GLS and GDH enzymatic activity in SIRT5 OE or KO cells with NH_4_Cl treatment was significantly enhanced by 1.27-fold and 1.33-fold or 1.20-fold and 1.08-fold (Figure S1). In accord, the results in MCN group were similar to those in SKN group. These results again verified that SIRT5 negatively regulates GLS and GDH enzymatic activity by lysine desuccinylation.

**Figure 7.**
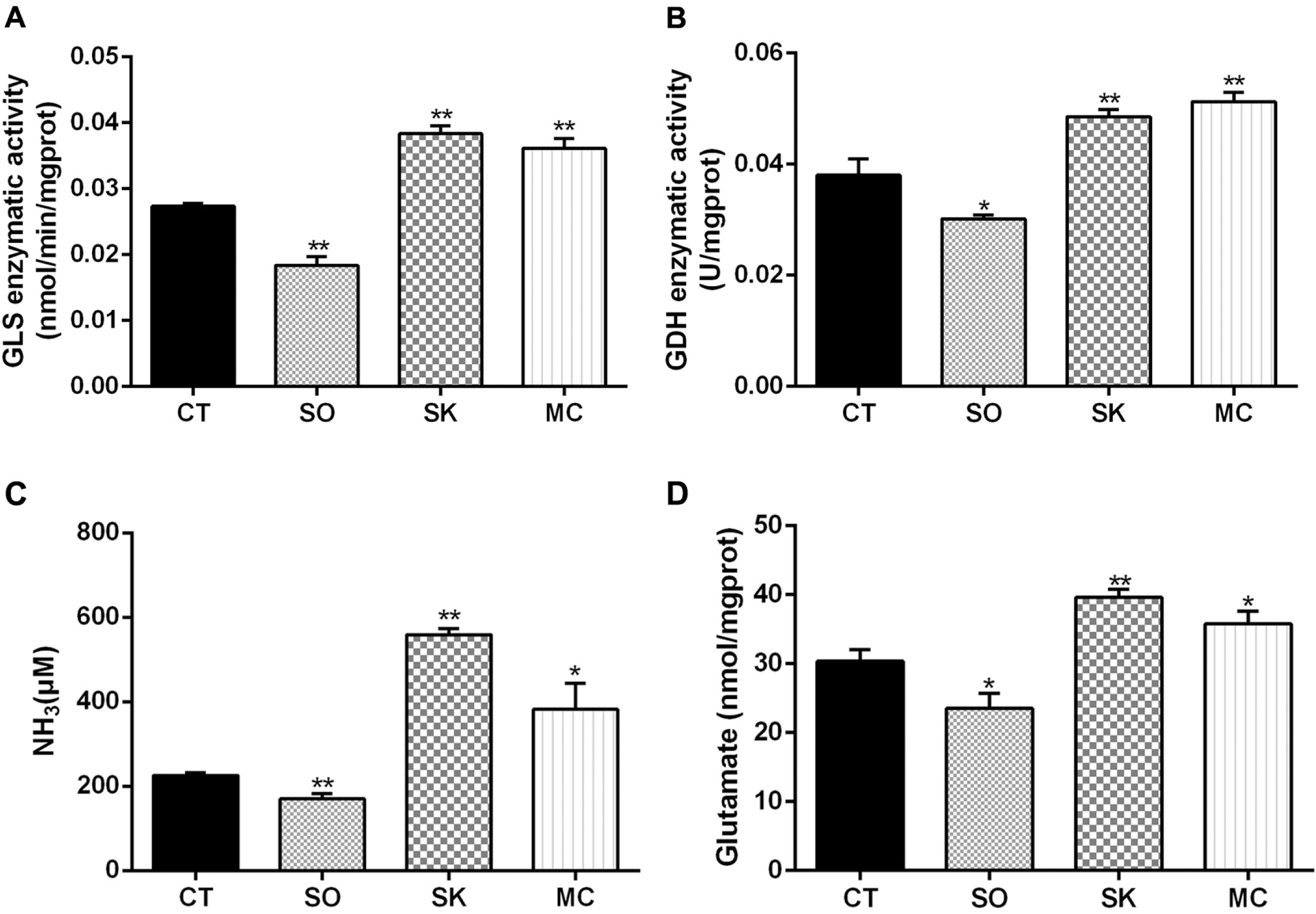
SIRT5 attenuated ammonia release. MAC-T cells (CT) were cultured in base medium. Cells in SIRT5 OE and SIRT5 KO group were cultured in base medium. MAC-T cells in MC group were treated by complete medium containing 20 μM MC3482 for 12 h. GLS activity in cells (A), GDH activity in cells (B), NH_3_ content in cells (C), Glutamate content in cells (D). Note: * P < 0.05, ** P < 0.01 vs .CT group.

**Figure 8.**
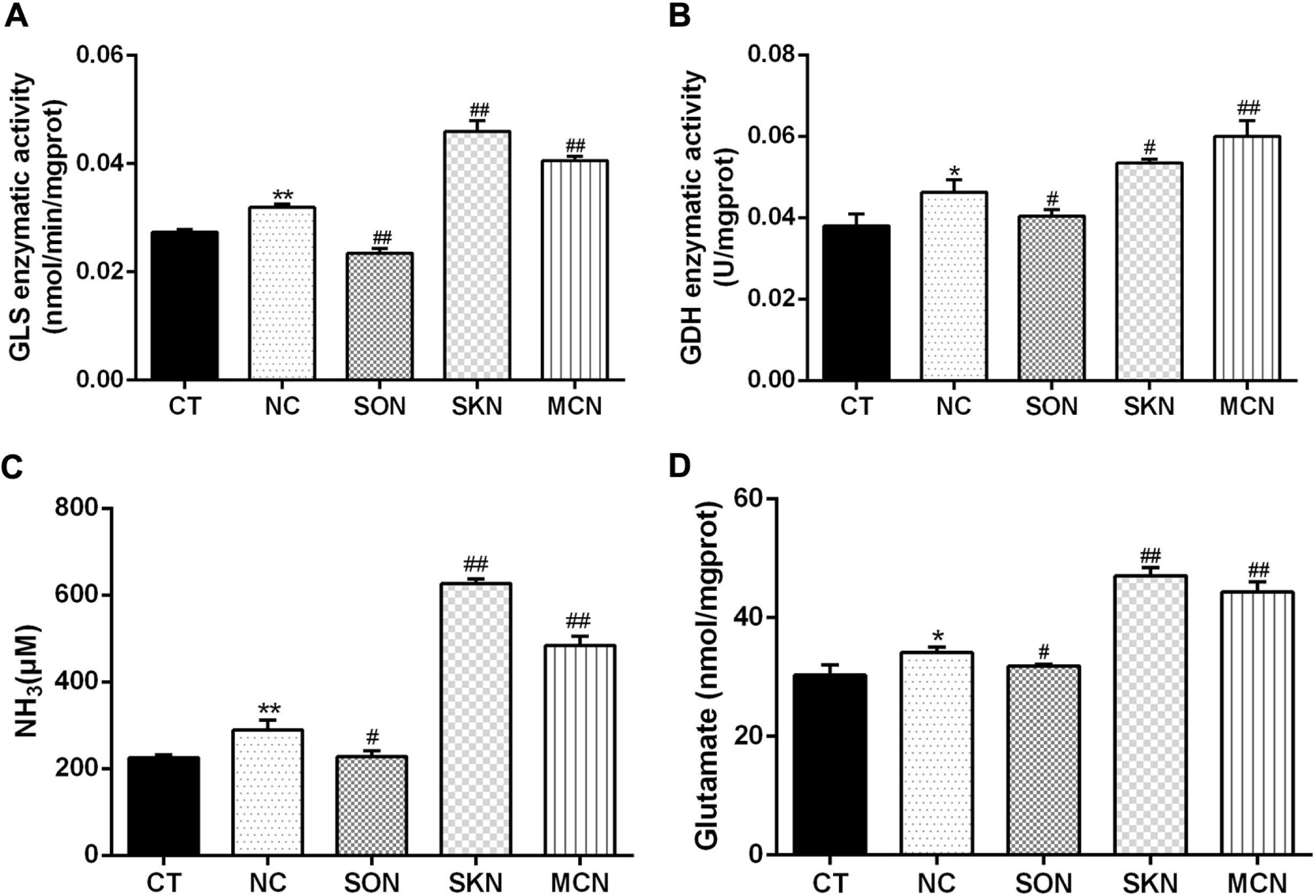
SIRT5 exacerbated ammonia release with NH_4_Cl treatment. MAC-T cells (CT) were cultured in base medium. MAC-T cells in NC group, and SO or SK cells in SON or SKN group were treated with base medium containing 10 mM NH_4_Cl for 12 h. MAC-T cells in MCN group were pretreated with 20 μM MC3482 for 30 min, and then treated with 10 mM NH_4_Cl for 12 h. GLS activity in cells (A), GDH activity in cells (B), NH_3_ content in cells (C), Glutamate content in cells (D). Note: * P < 0.05, ** P < 0.01 vs .CT group; # P <0.05, ## P < 0.01 vs. NC group.

We further used BPTES (GLS inhibitor) to treat cells, GLS enzymatic activity declined statistically significant, which suggested the significant inhibitory effect of BPTES on GLS enzymatic activity (Figure 9A). Moreover, the content of glutamate and NH_3_ were significantly down-regulated in MAC-T cells treated with BPTES (Figure 9B, 9C). As list in Figure 9B and 9C, compared with the BP group, the content NH_3_ and glutamate in SOBP group declined (P>0.05), while in the SKBP and MCBP groups, the content NH_3_ and glutamate increased (P>0.05). Altogether, SIRT5 inhibited glutamine metabolism in bovine mammary epithelial cells.

**Figure 9.**
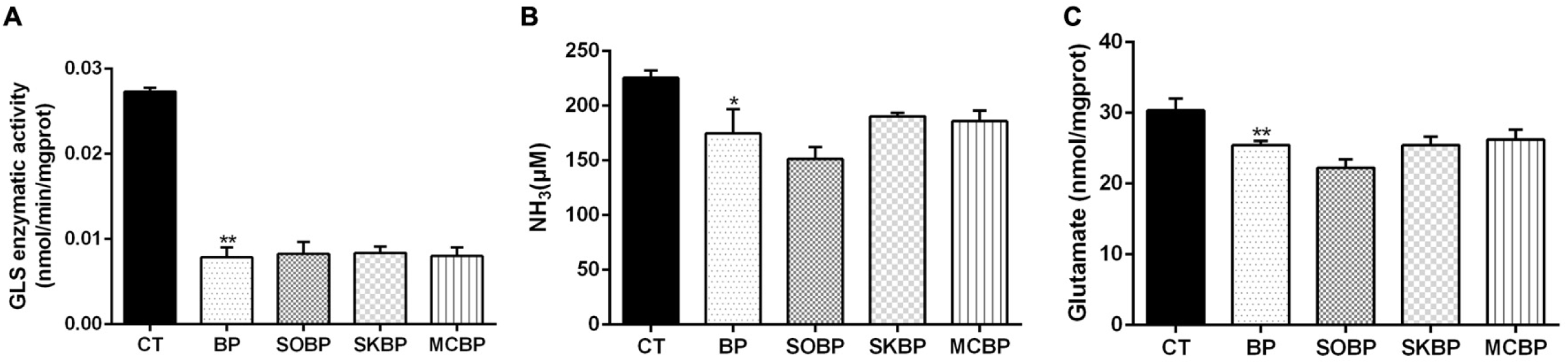
SIRT5 declined ammonia release with GLS inhibitor treatment. MAC-T cells (CT) were cultured in base medium. MAC-T cells in BP group, and SO or SK cells in SOBP or SKBP group were treated with base medium containing 0.12 μM BPTES for 12 h. MAC-T cells in MCBP group were pretreated with 20 μM MC3482 for 1 h, and then the cells were treated with 0.12 μM BPTES for 12 h. GLS activity in cells (A), NH_3_ content in cells (B), Glutamate content in cells (C). Note: * P < 0.05, ** P < 0.01 vs .CT group; # P <0.05, ## P < 0.01 vs. BP group.

### GLS and GDH were required for SIRT5 to regulate ammonia-induced cellular autophagy

Our previous studies found that SIRT5 reduced NH_3_-induced autophagy in bovine mammary epithelial cells [24]. We speculated that SIRT5 controlled the autophagy in bovine mammary epithelial cells by mediating GLS and GDH activity. As expected, compared to CT or NC group, the levels of Beclin1 and LC3II/I increased significantly and p62 expression decreased obviously in SIRT5 KO cells without or with NH_4_Cl treatment (Figure 10 and Figure 11). In contrast, the levels of Beclin1 and LC3II/I declined obviously and p62 expression increased significantly in SIRT5 OE cells without or with NH_4_Cl treatment (Figure 10 and Figure 11). Nevertheless, SIRT5 KO cells had more punctate LC3B positive cells and increased significantly autophagosome counts (Figure 12). After NH_4_Cl treatment, the number of punctate LC3B positive cells and autophagosomes in MAC-T cells and SIRT5 KO cells increased significantly, and the number of punctate LC3B positive cells and autophagosomes in SIRT5 KO cells were significantly more than those in the MAC-T cells (Figure 13). Generally, SIRT5 significantly reduces the number of dot shaped LC3B positive cells and autophagosomes, and inhibits the formation of autophagosomes. This was associated with elevated the autophagosomes number as well as increased protein levels of autophagic markers.

**Figure 10.**
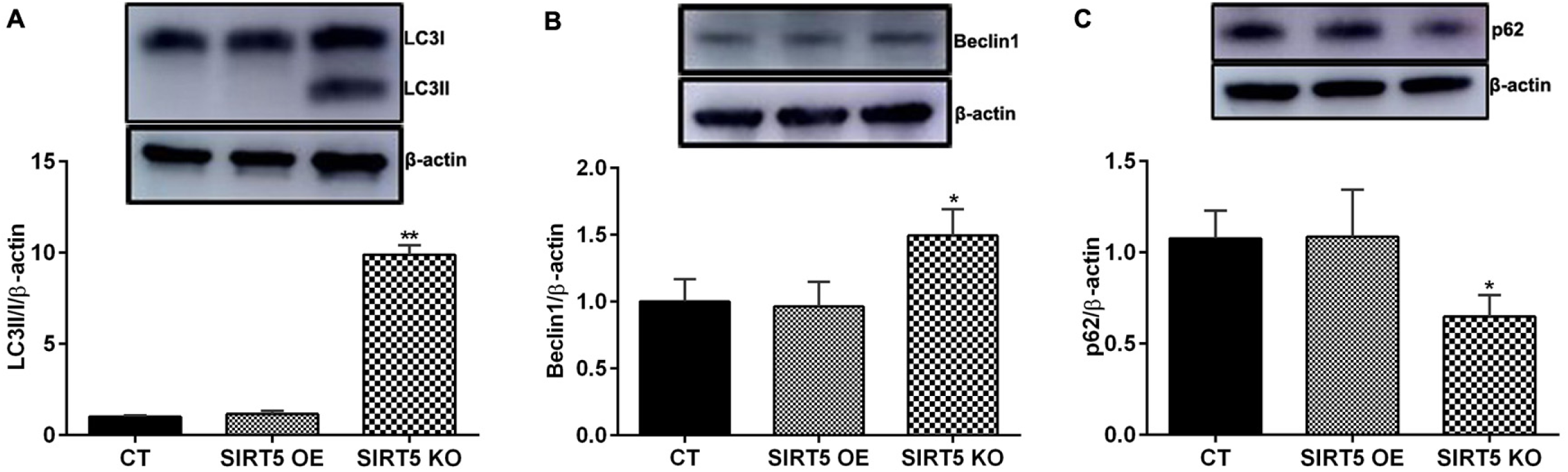
SIRT5 down-regulated the expression of Beclin1 and LC3II/I and increased p62 expression. MAC-T cells (CT), SO or SK cells were cultured in base medium. LC3II/I (A), Beclin1 (B), p62 (C). Note: * P < 0.05, ** P < 0.01 vs .CT group.

**Figure 11.**
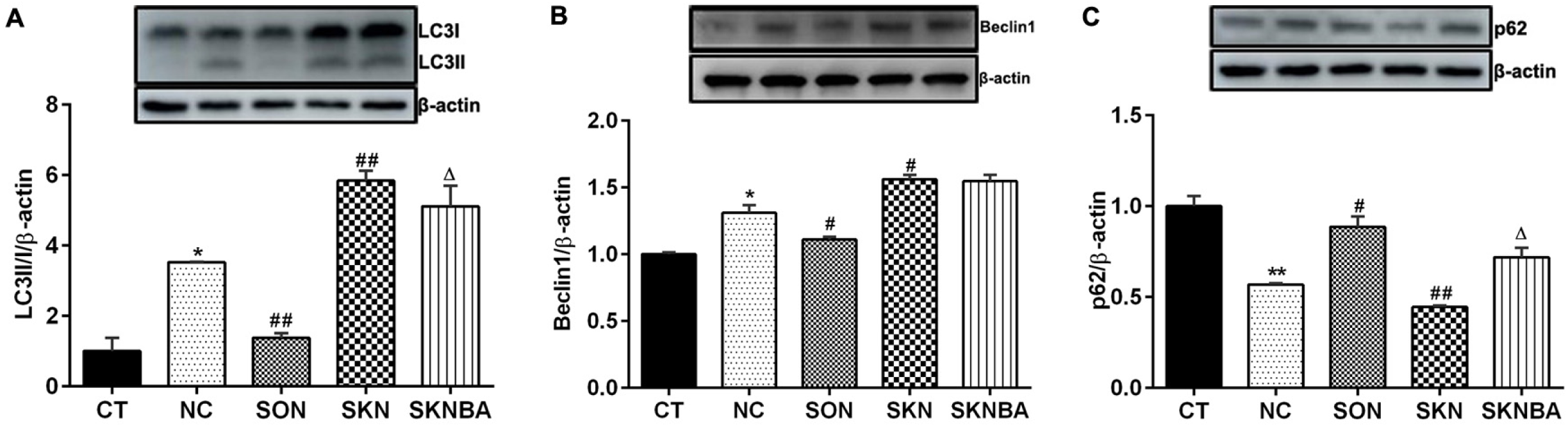
SIRT5 aggravated the decrease of Beclin1 and LC3II/I expression and the increase of p62 expression with NH_4_Cl treatment. MAC-T cells (CT) were cultured in base medium. MAC-T cells in NC group, and SO or SK cells in SON or SKN group were treated with base medium containing 10 mM NH_4_Cl for 12 h. In SKNBA group, SIRT5 KO cells were incubated with BafA1 (50 nM) for 1 h, and then the cells were treated with 10 mM NH_4_Cl for 12 h. LC3II/I (A), Beclin1 (B), p62 (C). Note: * P < 0.05, ** P < 0.01 vs .CT group; # P <0.05, ## P < 0.01 vs. NC group; Δ P < 0.05 vs .SKN group.

**Figure 12.**
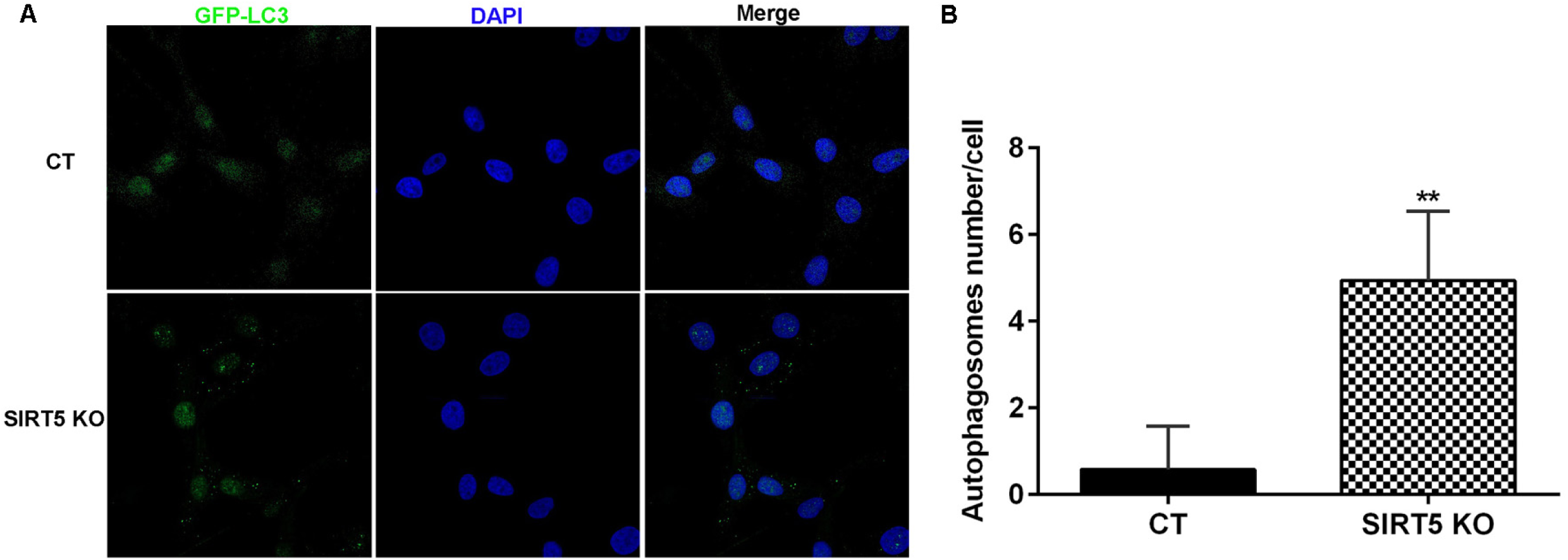
SIRT5 KO promoted the formation of autophagosomes. MAC-T cells (CT), and SK cells were cultured in base medium. GFP-LC3 staining (A), the number of autophagosomes (B). Note: ** P < 0.01 vs .CT group.

**Figure 13.**
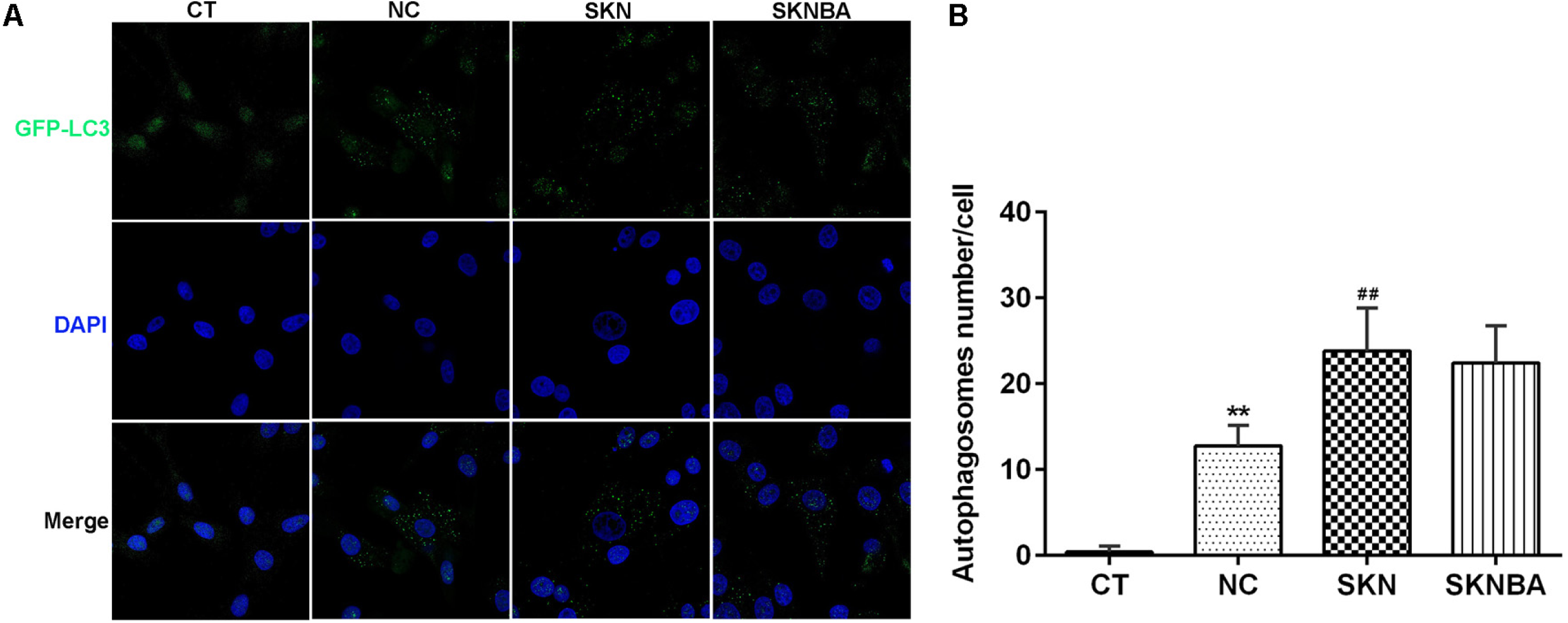
SIRT5 KO promoted the formation of autophagosomes with NH_4_Cl treatment. MAC-T cells (CT), and SK cells were cultured in base medium. MAC-T cells in NC group and SK cells in SKN group were treated with base medium containing 10 mM NH_4_Cl for 12 h. In SKNBA group, SIRT5 KO cells were incubated with BafA1 (50 nM) for 1 h, and then the cells were treated with 10 mM NH_4_Cl for 12 h. GFP-LC3 staining (A), the number of autophagosomes (B). Note: ** P < 0.01 vs .CT group; ## P < 0.01 vs. NC group; Δ P < 0.05 vs .SKN group.

Furthermore, we observed that SIRT5 KO led to NH_3_ accumulation, and this effect of SIRT5 KO was exacerbated by NH_4_Cl treatment (Figure S1 and Figure 11). This was associated with elevated autophagy as well as increased protein levels of autophagic markers (Figure 11). Additionally, we selected the inhibitor of autophagic flux (BafA1, 50 nM) to treat cells and found that BafA1 effectively diminished SIRT5 knockout-decreased p62 expression and elevated LC3II/I expression, had no effect on Beclin1 expression (Figure 11). There were similar changes in the number of autophagosomes (Figure 13). The above results suggested that SIRT5 interfered primarily with the autophagic flux, GLS and GDH were required as the major substrate of SIRT5. To sum up, these findings provided independent and in vitro data supporting that lysine succinylation could enhance GLS and GDH enzymatic activity, which interfered with ammonia-induced cellular autophagy.

### SIRT5 promoted the maintenance of mitochondrial homeostasis

SIRT5 in mitochondria decorated proteins, which were participated in metabolic activities, antioxidant pathways, energy production, apoptosis and autophagy, further maintaining mitochondrial homeostasis. Mitochondrial homeostasis involved in rebalancing the production and consumption of ATP during periods of negative energy balance [32–34]. Interestingly, our previous study found that SIRT5 declined the ratio of ADP/ATP [24]. In this study, we also found that SIRT5 increased the content of ATP (Figure 14A). Indeed, SIRT5 targeted succinate dehydrogenase complex, whose activity regulated the levels of intracellular reactive oxygen (ROS). Consistently, SIRT5 KO in mice increased ROS levels [35]. SIRT5 activated homeostatic mechanisms to protect cells from stressors [18]. Thus, we speculated that SIRT5 might affect autophagy by regulating mitochondrial homeostasis. As a result of SIRT5 OE cells had a greater number of mitochondria, with normal shapes and linear structures (Figure 14B). Compared with MAC-T cells, the number of mitochondria in SIRT5 OE cells was significantly increased, and the brightness was enhanced, indicating that SIRT5 OE increases the number and activity of mitochondria (Figure 14B). SIRT5 KO decreased the number of mitochondria, which were swelling and fragmentation (Figure 14B). Hence, SIRT5 had a role of maintaining mitochondrial homeostasis in bovine mammary epithelial cells.

**Figure 14.**
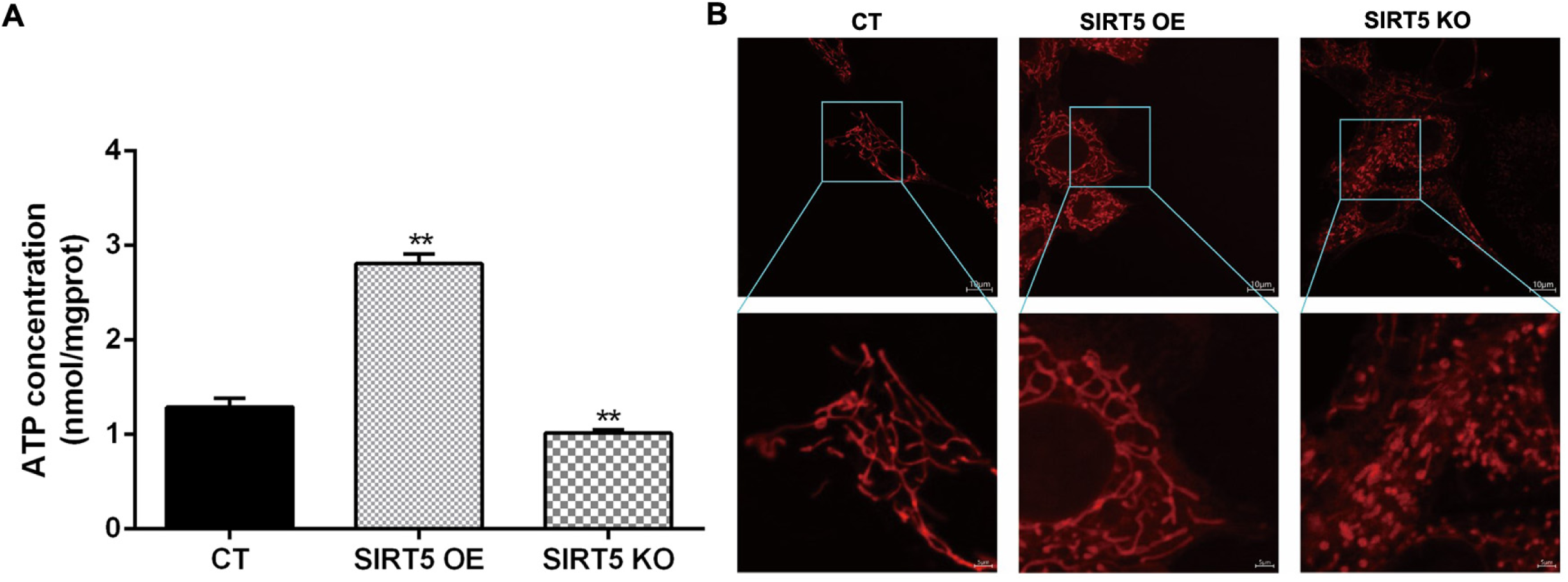
SIRT5 enhanced the content of ATP and the activity of mitochondria. MAC-T cells (CT), SO or SK cells were cultured in base medium. The content of ATP (A), Mito-Tracker Red-staining of mitochondria (B). Note: ** P < 0.01 vs .CT group.

## Discussion

Ammonia affected the health and growth of livestock, reducing their productivity [1]. NH_3_ increased the incidence rate and mortality of bovine respiratory diseases, and also reduced the productivity, lactation performance and pregnancy opportunities of cows [36, 37]. More than 98 % of ammonia in the body existed in the form of NH_4_^+^, which crossed the cell membrane through various transport channels and participated in various physiological activities [38]. In addition, NH_3_ could change pH, electrolyte balance and metabolic changes, which caused a negative impact on cellular function [2, 39]. High concentrations of NH_3_ could affect the composition of microorganisms in the nasal cavity and colon of rabbits, and interfere with local immune response and inflammatory processes, as well as increase the incidence of respiratory diseases in rabbits [40].

Autophagy exerted a significant role in maintaining the quantity and function of mammary epithelial cells, and was a key factor affecting the lactation performance of dairy cows. Ammonia had a dual role in autophagy, acting as an inducer at low concentrations and an inhibitor at high concentrations [3]. Our previous research found that NH_3_ regulated autophagy of cow mammary epithelial cells through the PI3K/Akt/mTOR signaling pathway [14]. Interestingly, SIRT5 was a key regulatory factor in maintaining cellular homeostasis, which regulated protein activity in various metabolic processes with post-translational modifications and controlled ammonia induced autophagy in tumor cells [15–22]. In addition, we found that SIRT5 inhibited autophagy of cow mammary epithelial cells, and the PI3K/Akt/mTOR signaling was involved in NH_3_ induced autophagy, depending on SIRT5 [24]. Meanwhile, SIRT5 promoted autophagy in colorectal cancer, and its overexpression was associated with low survival rates [20]. Moreover, the AMPK/mTOR pathway was involved in SIRT5-enhanced autophagy in gastric cancer cells, while SIRT5 expression was suppressed in gastric cancer tissues [25]. Strangely, in breast cancer and mouse myogenic cells, silencing SIRT5 increased ammonia-induced autophagy by controlling glutamine metabolism and mitophagy [12]. However, SIRT5 had been shown to exhibit weaker deacetylase activity, more commonly demalonylase, desuccinylase, and deglutarylase activities, participating in several metabolic processes of mitochondria [18, 28–31]. SIRT5 downregulated the succinylation level of GLS and inhibited ammonia-induced autophagy in tumor cells [12]. Our previous studies also found that SIRT5 reduced NH_3_-induced autophagy in bovine mammary epithelial cells, but little is known about the specific molecular mechanism.

Bcl-2 associated athanogene 3 (BAG3) stabilized GLS through inhibiting its interaction with SIRT5, thereby preventing its desuccinylation to enhance autophagy [41]. SIRT5 regulated GDH to provide glutamine with entering the tricarboxylic acid (TCA) cycle in malignant colorectal cancer cells [42]. Mechanistically, the direct interaction between SIRT5 and GDH led to deglutarylation and functional activation of GDH, which was a key regulatory factor of glutaminolysis [42]. Interestingly, our recent study found that SIRT5 reduced the content of NH_3_ and glutamate in bovine mammary epithelial cells by inhibiting GLS activity, and declined the ratio of ADP/ATP [24]. Therefore, we hypothesized that SIRT5-mediated GLS and GDH desuccinylation controlled ammonia production, further regulated the autophagy of mammary epithelial cells. To verify this point, we constructed SIRT5 overexpression or knockout cell lines. Additionally, ammonium chloride, and some inhibitors for SIRT5, GLS, and autophagy were also used to treat cells. Immunoprecipitation techniques verified that SIRT5 interacted with endogenous GLS or GDH. Strangely, SIRT5 had no effect on the expression of endogenous GLS or GDH.

Reversible post-translational modifications are considered as key regulators of mitochondrial proteins and various metabolism. The analysis revealed the potential impact of lysine succinylation on enzymes of mitochondrial metabolism, such as tricarboxylic acid cycle (TCA), fatty acid metabolism, and amino acid degradation [43]. SIRT5 with efficient lysine desuccinylase inhibited the biochemical activity of pyruvate dehydrogenase complex and succinate dehydrogenase [43]. SIRT5 deficient in mice appeared to enhance succinylation level of carbamoyl phosphate synthase 1, which was a known target of SIRT5 [29]. The absence of SIRT5 caused the accumulation of medium-chain and long-chain acylcarnitines, and reduced the production of β-hydroxybutyric acid [23]. Moreover, SIRT5 mediated succinylation level of ketogenic enzyme 3-hydroxy-3-methylglutaryl-CoA synthase 2 (HMGCS2) in vivo and in vitro [23]. In summary, SIRT5 was a global regulator for mitochondrial lysine succinylation. Interestingly, our previous study found that SIRT5 reduced the content of NH_3_ and glutamate in bovine mammary epithelial cells by inhibiting GLS activity [24]. Based on SIRT5, which had a strong desuccinylation effect, we determined the protein succinylation level and demonstrated that SIRT5 reduced the protein succinylation level in cells. Interestingly, we also found that SIRT5 attenuated the succinylation levels of GLS and GDH, which were consistent with the results of total protein.

To determine the functional consequence of SIRT5 catalyzing the desuccinylation of GLS and GDH lysine, we examined GLS and GDH enzymatic activity, and demonstrated that SIRT5 reduced the enzymatic activity of GLS and GDH, as well as the content of NH_3_ and glutamate. Correspondingly, after treatment with NH_4_Cl, SIRT5 also reduced GLS and GDH enzymatic activity, resulting in the induction of NH_3_ and glutamate content and alleviating glutamine metabolism. Meanwhile, inhibiting SIRT5 (MC3482, a specific SIRT5 inhibitor) improved obviously GLS and GDH enzymatic activity, accompanying with obvious reduction in the content NH_3_ and glutamate. These results again verified that SIRT5 regulated negatively GLS and GDH enzymatic activity by lysine desuccinylation. We further treated cells with BPTES (GLS inhibitor) and confirmed that SIRT5 inhibited glutamine metabolism in bovine mammary epithelial cells. Collectively, we found that SIRT5 obviously decreased the succinylation levels of GLS and GDH, and further decreased the enzymatic activity of GLS and GDH.

Autophagy was a strictly mediated process that got rid of damaged organelles or cytosolic components, the process of which started with the formation of autophagosomes with double-membrane vesicles [44]. Then, the melt autophagosomes with cargo were transported to lysosomes for degradation and recovery. LC3, Beclin1 and p62 are indicated genes known as autophagy related genes. Moreover, autophagy was also affected by many other genes [45]. There is evidence that SIRT5 had a regulatory function in autophagy [25, 46]. SIRT5 promoted autophagy in colorectal cancer and gastric cancer cells [20, 25]. In breast cancer and mouse myogenic cells, silencing SIRT5 increased ammonia-induced autophagy by controlling glutamine metabolism and mitophagy [12]. Another study showed that SIRT5 increased autophagy and accelerated in colorectal cancer growth by deacetylating lactate dehydrogenase B, thereby promoting its activity, and knockdown or inhibition of SIRT5 down-regulated autophagy levels and inhibited growth [47].

There was much evidence that both ammonia and SIRT5 could regulate the level of cellular autophagy, and excess ammonia increased the expression of the classical autophagy genes LC3 and Beclin1, and decreased the expression of p62/SQSTM1, suggesting that high ammonia induced autophagy of skeletal muscle in patients with cirrhosis [48], and that ammonia promoted the occurrence of autophagy in hepatocytes in mice with high blood ammonia [49]. Our previous studies also found that SIRT5 reduced NH_3_-induced autophagy in bovine mammary epithelial cells [24]. It was known that GLS and GDH participate in the regulation of ammonia production [2, 50]. Therefore, we speculated that SIRT5 regulated GLS and GDH activity to control the autophagy of bovine mammary epithelial cells. As expected, SIRT5 attenuated the levels of Beclin1 and LC3II/I, and enhanced p62 expression without or with NH_4_Cl treatment. Nevertheless, SIRT5 significantly reduced the number of dot shaped LC3B positive cells and autophagosomes, and inhibited the formation of autophagosomes. This was associated with elevated the autophagosomes number as well as increased protein levels of autophagic markers. Furthermore, we observed that SIRT5 KO led to NH_3_ accumulation, and this effect of SIRT5 KO was exacerbated by NH_4_Cl treatment. This was associated with elevated autophagy as well as increased protein levels of autophagic markers (Figure 11). Additionally, we selected the inhibitor of autophagic flux (BafA1) to treat cells and found that BafA1 effectively diminished SIRT5 knockout-decreased p62 expression and elevated LC3II/I expression, had no effect on Beclin1 expression. There were similar changes in the number of autophagosomes (Figure 13). Hence, these findings provided independent and in vitro data supporting that lysine succinylation could enhance GLS and GDH enzymatic activity, which influenced ammonia-induced cellular autophagy. These results suggested that SIRT5 interfered primarily with the autophagic flux, GLS and GDH were required as the major substrate of SIRT5.

Mitochondria are the central organelles of metabolic activities, and several factors have been demonstrated to maintain these homeostatic mechanisms [51, 52]. Due to the important role of sirtuins in regulating metabolism, this protein family has aroused great interest in the scientific community. Especially, SIRT3, SIRT4, and SIRT5 located in the mitochondrial matrix, where they mediated proteins participating in metabolic responses, energy metabolism, antioxidant roles, and autophagy, further maintaining the homeostasis of mitochondria [32, 33]. SIRT5 had been described to play a role in mediating cellular metabolism, detoxification, oxidative stress, energy balance, and autophagy [53]. Pathway analysis indicated that SIRT5 targeted fatty acid β-oxidation, branched chain amino acid catabolism, the citric acid cycle, ATP synthesis, and ketone body synthesis, which was the most enriched target pathways [23]. The activity and expression of mitochondrial sirtuins were closely related to cellular metabolic status, making metabolic adaptation possible, thereby rebalancing ATP production and consumption during negative energy balance periods [34]. Interestingly, our previous study found that SIRT5 declined the ratio of ADP/ATP [24]. In this study, we also found that SIRT5 increased the content of ATP (Figure 14A). Indeed, SIRT5 targeted succinate dehydrogenase complex, whose activity could interfere with the intracellular reactive oxygen (ROS) levels [35]. SIRT5 maintained mitochondrial homeostasis to protect cells from stressors [18]. Thus, we speculated that SIRT5 might affect autophagy by regulating mitochondrial homeostasis. As a result of SIRT5 OE cells had a greater number of mitochondria, with normal shapes and linear structures (Figure 14B). Compared with the MAC-T cells, the number of mitochondria in SIRT5 OE cells was significantly increased, and the brightness was enhanced, indicating that SIRT5 OE increases the number and activity of mitochondria (Figure 14B). SIRT5 KO decreased the number of mitochondria, which were swelling and fragmentation (Figure 14B). Hence, SIRT5 had a role of maintaining mitochondrial homeostasis in bovine mammary epithelial cells.

Collectively, we concluded that SIRT5 reduced ammonia release by modulating the succinylation levels and enzymatic activities of GLS and GDH in mitochondria and promoted the maintenance of mitochondrial homeostasis, as well as further attenuated ammonia-induced autophagy in bovine mammary epithelial cells (Figure 15). Consistently, we discovered that GLS and GDH were two physiological substrates of SIRT5, which relied on desuccinylation to reduce ammonia production by inhibiting the enzymatic activity of GLS and GDH.

**Figure 15.**
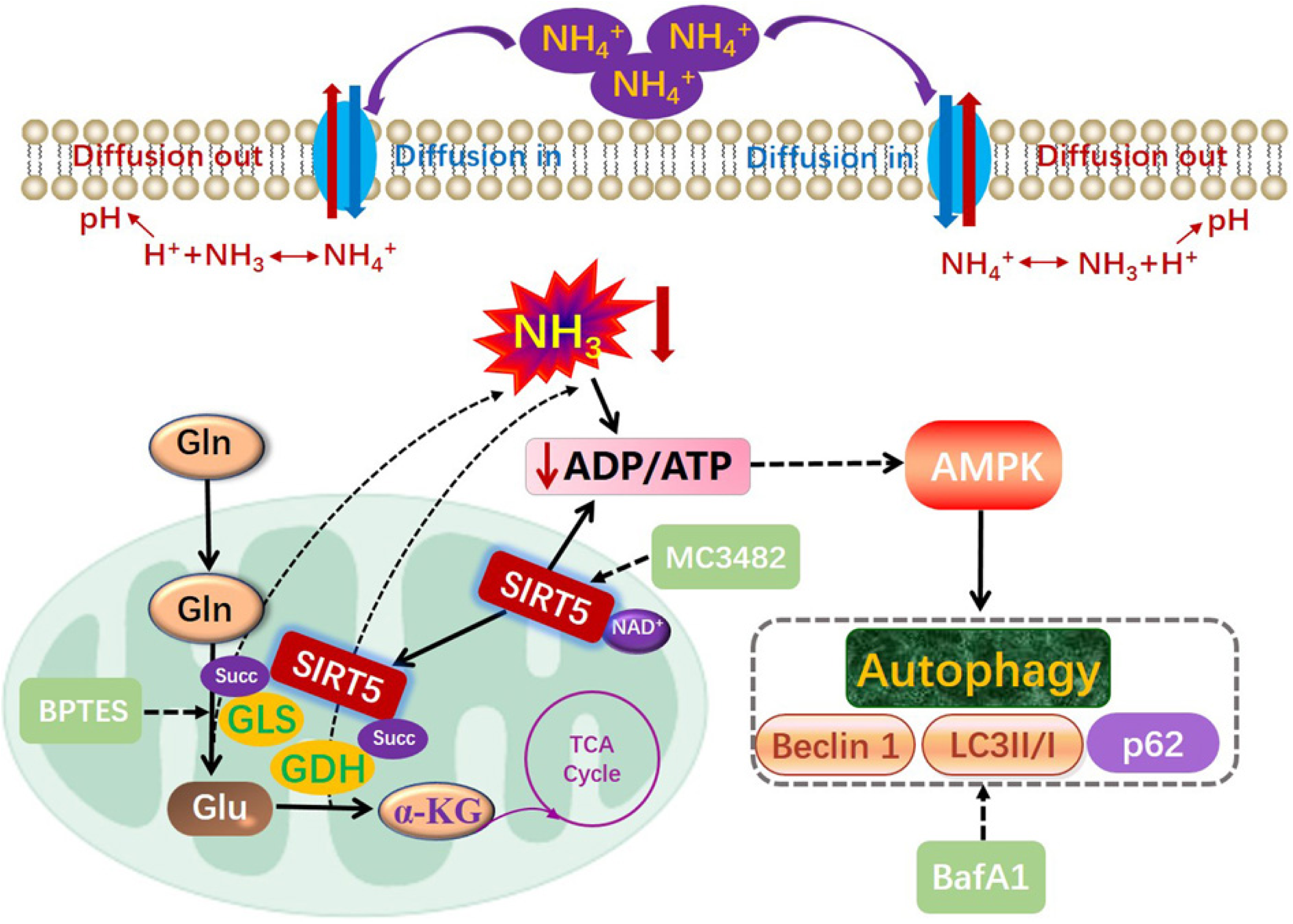
A model of SIRT5-mediated GLS and GDH desuccinylation attenuated the autophagy of bovine mammary epithelial cells induced by ammonia. GLS and GDH were two physiological substrates of SIRT5, which relied on desuccinylation to reduce ammonia production by inhibiting the enzymatic activity of GLS and GDH in mitochondria. SIRT5 enhanced ATP content and promoted the maintenance of mitochondrial homeostasis, as well as further attenuated ammonia-induced autophagy in bovine mammary epithelial cells. SIRT5, sirtuin 5; LC3B, light chain 3 β; GLS, glutaminase; α-KG, α-ketoglutarate; Gln, glutamin; GDH, glutamate dehydrogenase; Glu, glutamate; BPTES, GLS inhibitor; MC3482, SIRT5 inhibitor; BafA1, bafilomycin A1, autophagy inhibitor. The solid arrow represented the strengthening effect. The dashed arrow represented the inhibitory effect.

## Availability of data and materials

The datasets used and/or analyzed during the present study are available from the corresponding author upon request.

## Author contributions

LHP and WYY conceived and designed the experiment. YHL, GSK, LGY and JJH executed experiments; DJR, ZXY and LLY cultured and treated cells; ZK and GS drew figures, ZGM and HLQ performed data analyses; YHL, GSK, LHP, and WYY interpreted the data and wrote the manuscript.

## Ethics approval and consent to participate

This article does not contain any studies with human subjects or animals performed by any of the authors. All experimental protocols were approved by College of Animal Veterinary Medicine, Henan Agricultural University Ethics Committee (Zhengzhou, China).

## Patient consent for publication

Not applicable.

---

**Figure S1.**
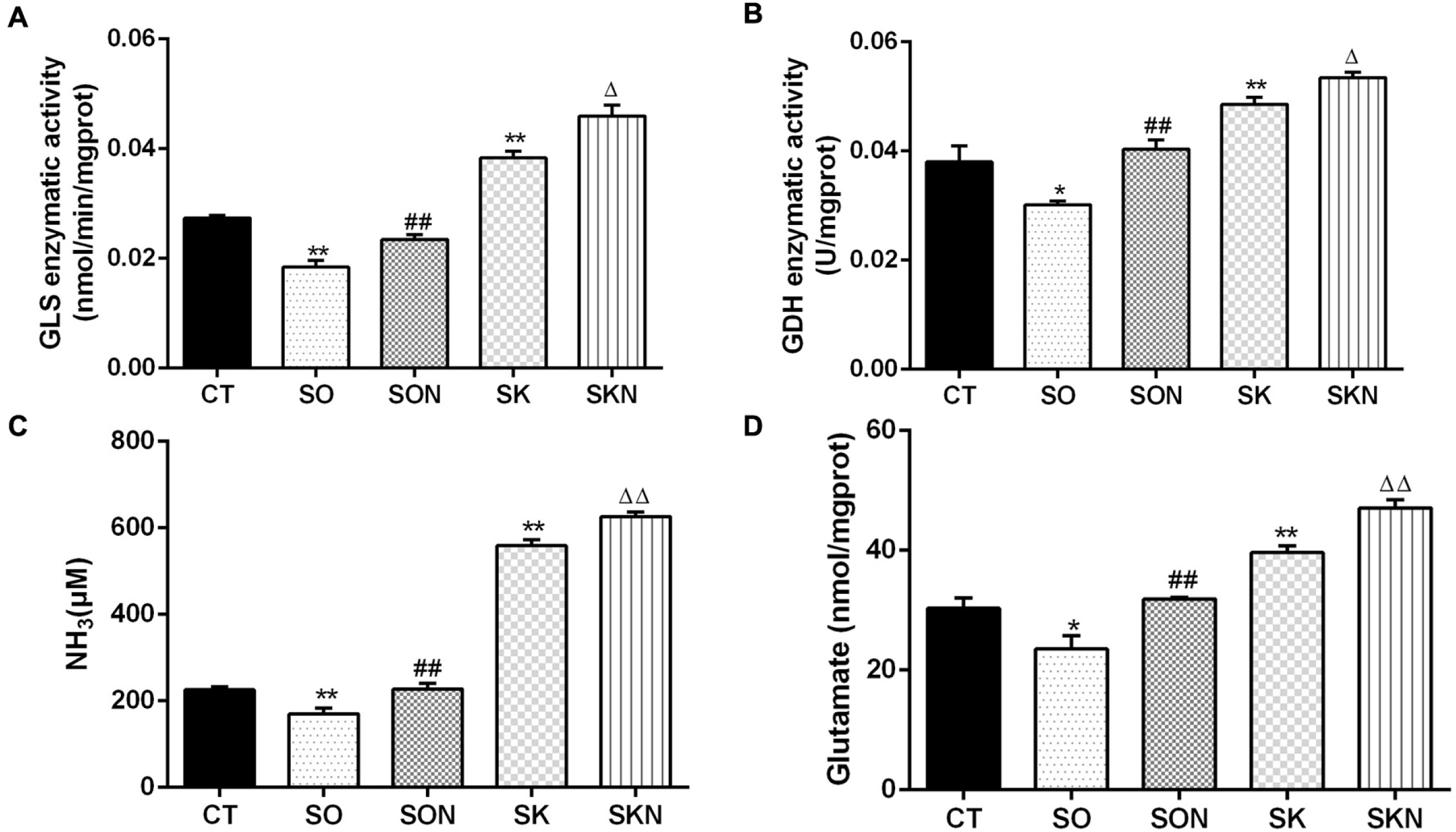
NH_4_Cl treatment exacerbated the effect of SIRT5 on ammonia release. MAC-T cells (CT), SO or SK cells were cultured in base medium. SO or SK cells in SON or SKN group were treated with base medium containing 10 mM NH_4_Cl for 12 h. GLS activity in cells (A), GDH activity in cells (B), NH_3_ content in cells (C), Glutamate content in cells (D). Note: * P < 0.05, ** P < 0.01 vs .CT group; ## P < 0.01 vs. SO group; Δ P < 0.05, ΔΔ P < 0.01 vs .SK group.

